# The tri-culture system reveals an activation cascade from microglia through astrocytes to neurons during neuroinflammation

**DOI:** 10.1101/2025.07.27.667064

**Authors:** Hayato Kobayashi, Hiroshi Kato, Mitsuho Taniguchi, Setsu Endoh-Yamagami

## Abstract

Neuroinflammation is involved in various neurodegenerative diseases, with glial cells playing crucial roles. It is known that neuroinflammation is initiated by microglia, which interact with astrocytes and neurons. However, the detailed molecular mechanisms underlying intercellular interactions during neuroinflammation are not fully understood. To elucidate these mechanisms, multicellular culture systems are required, although the availability of the culture systems using human cells is limited. In this study, we developed a tri-culture system of neurons, astrocytes, and microglia derived from human induced pluripotent stem cells (iPSCs) to evaluate their relationships in neuroinflammation. Previously, differentiation of astrocytes from iPSCs was commonly induced using serum. However, serum stimulation has been reported to cause irreversible activation of astrocytes. Therefore, we generated astrocytes using a serum-free method and established a tri-culture system. Microglia cocultured with the astrocytes and neurons exhibited a morphology with branched processes compared to the monoculture system, suggesting a homeostatic state. By applying lipopolysaccharide (LPS) stimulation to induce inflammation, the microglial morphology shifted to an amoeboid shape, accompanied by an increase in the expression of pro-inflammatory cytokines. Additionally, nuclear translocation of NF-κB revealed that LPS specifically activates microglia through the TLR4 receptor, which subsequently releases TNF-α, leading to the activation of astrocytes. Furthermore, activated astrocytes were shown to enhance neuronal excitability. Using the tri-culture system, we elucidated a part of the cascade involving microglia, astrocytes, and neurons during neuroinflammation and demonstrated the amplification of inflammatory signals through cell communication. This culture system will be valuable for conducting detailed investigations into the interactions between glia and neurons, advancing research on neurodegenerative diseases associated with neuroinflammation.

## 1 Introduction

Microglia are immune cells in the brain, serving a vital role in defending the brain against pathogens and tissue damages. Microglia quickly detect changes in the brain environment, and they remove pathogens and cellular debris, subsequently triggering inflammation (Xu et al., 2016). Microglia are characterized by a branched morphology in the homeostatic state, but upon activation, they undergo a morphological transformation into an amoeboid shape without branched processes, accompanied by the release of pro-inflammatory cytokines (Vidal-Itriago et al., 2022). Although neuroinflammation is a defensive response of the brain, chronic neuroinflammation is known to have detrimental effects on the brain, contributing to neurodegenerative diseases, psychiatric disorders, and chronic pain (Kwon et al., 2020; Brites et al., 2015; Ji et al., 2014). In a neuroinflammatory state, it has been reported that microglia interact with astrocytes through pro-inflammatory cytokines. Activated microglia stimulate astrocytes through TNF-α (Guttikonda et al., 2021). Astrocytes stimulated by activated microglia transition into a state called A1 astrocytes, which can cause damage to neurons and are involved in various neurodegenerative diseases (Liddelow et al., 2017). Glial cells are also known to exert an influence on neuronal function (Cserép et al., 2020; Linnerbauer et al., 2020). Previous research using rodents has reported that LPS treatment induces neuroinflammation, resulting in enhanced neuronal excitability and seizures (Rodgers et al., 2009). However, there is a lack of in vitro models that can evaluate the activation of glial cells and subsequent changes in neuronal activity, limiting the detailed mechanistic analysis of neuroinflammation.

It is known that glial cells exhibit species differences (Zhang et al., 2016; Geirsdottir et al., 2019), and it is important to utilize human-derived glial cells when studying neuroinflammation in humans. Previously, it was difficult to acquire human brain-derived cells. However, the invention of induced pluripotent stem cells (iPSCs) has made it relatively easy to obtain brain cells (Abud et al., 2017; Tchieu et al., 2019; Badanjak K et al., 2021; Stöberl et al., 2023). Nevertheless, there are several challenges in studies using human iPSC-derived neural cells and microglia. Firstly, the monoculture of iPSC-derived microglia has been reported to exhibit an amoeboid morphology and result in microglial activation even without stimulation (Baxter et al., 2021). Additionally, the differentiation method using serum has been commonly used for generating astrocytes from iPSCs. However, it has been reported that astrocytes exposed to serum undergo irreversible activation (Foo et al., 2011).

In this study, we constructed a tri-culture system using human iPSC-derived neurons, astrocytes, and microglia without exposing them to serum. The tri-culture system successfully reproduced microglia with branched processes, which represents the homeostatic state. The tri-culture system was utilized to induce neuroinflammation through application of lipopolysaccharide (LPS), a known microglia stimulant. In the tri-culture system, the subcellular localization of nuclear factor kappa B (NF-κB), an important protein involved in the inflammatory response, was evaluated to visualize the temporal cell-to-cell interactions. Furthermore, we demonstrated that neuronal activity was increased in the presence of astrocytes activated by tumor necrosis factor-alpha (TNF-α) released from microglia. This tri-culture system will be useful for uncovering the pathogenesis of neuroinflammation-related diseases.

## 2 Materials an methods

### 2.1 Generation of astrocytes from iPSC-derived astrocyte progenitor cells

Astrocytes were generated from iPSC-derived astrocyte progenitor cells (XCell Science). The astrocyte progenitor cells were maintained in the progenitor medium containing 1% N-2 MAX Media Supplement, 1% Glutamax, 1% MEM non-essential amino acids solution, 1% penicillin-streptomycin solution, 0.5 U/mL heparin, 10 ng/mL bFGF, and 10 ng/mL EGF in DMEM/F12. To differentiate the astrocyte progenitor cells into astrocytes, the cells were initially plated in the progenitor medium. The medium was then changed to the differentiation medium containing 1% N-2 MAX Media Supplement, 1% Glutamax, 1% MEM non-essential amino acids solution, 1% penicillin-streptomycin solution, 0.5 U/mL heparin, 10 ng/mL CNTF, and 10 ng/mL BMP4 in DMEM/F12 on the following day. After 5 days of differentiation, astrocytes were used for gene expression analysis and immunostaining.

### 2.2 qRT-PCR

RNA extraction was performed using the FastLane Cell SYBR Green Kit (QIAGEN) or QIAseq UPX Cell Lysis Kit (QIAGEN). Quantitative mRNA expression analysis was performed using the QuantiTect SYBR Green RT-PCR Kit (QIAGEN) according to the manufacture’s protocol. The following primers were used: *GAPDH*, sense: 5’-gtcagtggtggacctgacct-3’, antisense: 5’-tgctgtagccaaattcgttg-3’; *S100B*, sense: 5’-tggccctcatcgacgttttc-3’, antisense: 5’-atgttcaaagaactcgtggca-3’; *KCNJ10*, sense: 5’-cgcacagctctgctcctaac-3’, antisense: 5’-ccggctttctgtctgagtgg-3’; *AQP4*, sense: 5’-gagtgacagacccacagcaa-3’, antisense: 5’-catggccagaaattccgctg-3’; *SLC1A2*, sense: 5’-gattcgtccttcctgttgga-3’, antisense: 5’-agcaggctgatgtcctctgt-3’; *SLC1A3*, sense: 5’-gagtcaatgccctgggtcta-3’, antisense: 5’-gtcacggtgtacatggcaag-3’; *GFAP*, sense: 5’-ctgcggctcgatcaactca-3’, antisense: 5’-tccagcgactcaatcttcctc-3’; *TNF*, sense: 5’-ctcttctgcctgctgcactttg-3’, antisense: 5’-atgggctacaggcttgtcactc-3’; *IL1B*, sense: 5’-cgaggcacaaggcacaacag-3’, antisense: 5’-agggccatcagcttcaaagaac-3’. PCR was conducted using the CFX384 Touch Real-Time PCR Detection System (Bio-Rad). The data were analyzed using the delta-delta Ct method using GAPDH as the endogenous control.

### 2.3 Immunofluorescence

The cultured cells were fixed with 4% formaldehyde solution and washed three times with PBS (−). Subsequently, blocking and permeabilization were performed using PBS (−) supplemented with 1% BSA and 0.2% Triton X-100 for 1 hr. Then, the cells were incubated overnight at 4°C with primary antibody solution (anti-IBA1, rabbit, 1:800, Fujifilm Wako; anti-MAP2, chicken, 1:800, Novus Biologicals; anti-GFAP, mouse, 1:3000, Merck Millipore; anti-IBA1, goat, 1:500, Fujifilm Wako; anti-NF-κB, rabbit, 1:400, Cell Signaling; anti-NeuN, mouse, 1:700, Sigma Aldrich). After three times of PBS (−) wash, cells were incubated with secondary antibodies and Hoechst 33342 (Dojindo) for 1 hr at room temperature (anti-rabbit Alexa Fluor 488, goat, 1:1000, Thermo Fisher Scientific; anti-chicken Alexa Fluor 594, goat, 1:1000, Thermo Fisher Scientific; anti-mouse Alexa Fluor 647, goat, 1:1000, Thermo Fisher Scientific; anti-goat Alexa Fluor 488, donkey, 1:1000, Thermo Fisher Scientific; anti-rabbit Alexa Fluor 594, donkey, 1:1000, Thermo Fisher Scientific). After three times wash with PBS (−), images were captured using ECLIPSE Ti (Nikon).

### 2.4 Monoculture of iCell Microglia

iCell Microglia (Fujifilm Cellular Dynamics) was used as human iPSC-derived microglia. The cells were cultured according to the application protocol provided by Fujifilm Cellular Dynamics. Briefly, the cells were suspended in iCell Microglia Maintenance Medium and seeded onto 96-well plate coated with 0.07% polyethyleneimine (3×10^4^ cells/well). The cells were cultured at 37°C with 5% CO2. The medium was changed with iCell Microglia Maintenance Medium every 3-4 days. After 28 days, the cells were fixed using formaldehyde solution.

### 2.5 Generation of iPSC-derived Neurons

iPSC-derived neurons were generated by enforced expression of the *Ngn2* (Wang et al., 2017). The enforced expression of *Ngn2* was achieved by inserting the doxycycline-inducible mouse *Ngn2* gene into the genome using the AAVS1 Safe Harbor Targeting System (System Biosciences) as described previously (Kobayashi et al., 2024). The iN-iPSCs were cultured in StemFit medium until they reached approximately 80% confluency. Subsequently, the cells were collected using TrypLE™ Select Enzyme and seeded onto a Matrigel coating plate at a density of 1×10^5^ cells/cm^2^ using the neuronal differentiation medium containing 0.5% B-27 supplement minus vitamin A, 1% Glutamax, 1% penicillin-streptomycin solution, 2 μg/mL doxycycline, and 5 μM Y-27632 in Neurobasal medium. Medium changes were performed after 1 and 4 days of passage, and after 6 days of passage, the iPSC-derived neurons were cryopreserved.

### 2.6 Generation of neuron/astrocyte/microglia tri-culture system

The astrocyte progenitor cells were seeded onto 96-well plates at a density of 6×10^4^ cells/well using the progenitor medium. After 1 day, the astrocyte progenitor cells were differentiated into astrocytes by replacing with the differentiation medium. After 5 days of differentiation, iPSC-derived neurons and iCell microglia were seeded onto the astrocytes at a density of 3×10^4^ cells/well each. The medium was changed every 1-3 days. After 3 to 4 weeks culture, the cells were stimulated with 100 ng/mL of LPS. TNF-α antagonist and TLR4 inhibitor were added simultaneously with LPS stimulation. After 0, 0.5, 1, 3, 6, 24 hours of LPS stimulation, the cells were fixed using a formaldehyde solution.

### 2.7 Measurement of TNF-α and IL-1β proteins

The microglia monoculture system, cultured in the differentiation medium, and the tri-culture system were both stimulated with LPS at a final concentration of 100 ng/mL for 24 hours. Following stimulation, the culture medium was collected and centrifuged at 2,000 × g for 5 minutes at room temperature. Subsequently, the supernatants were analyzed for the levels of TNF-α and IL-1β. Cytokine quantification was conducted using the V-PLEX Human Proinflammatory Panel II (4-Plex) kit (Meso Scale Discovery, MSD), following the manufacturer’s protocol. Samples were diluted 10-fold with Diluent 2 prior to measurement. Signal detection was performed using the Meso QuickPlex SQ 120 MM instrument (MSD).

### 2.8 Ca^2+^ imaging assay

Four-week co-cultured cells were used for the Ca^2+^ imaging assay. The cells were incubated for 1 hr at 37°C with 50μL of culture medium and 50 μL of working solution containing Calbryte 520 AM (AAT Bioquest), 2.5 mM calcium chloride dihydrate (Fujifilm Wako), 0.04% pluronic F-127 (Thermo Fisher Scientific), and 10 mM HEPES buffer solution (Fujifilm Wako) in HBSS (–) without phenol red (Fujifilm Wako). Then the medium was replaced with a recording solution containing 2.5 mM calcium chloride dihydrate, 10 mM HEPES buffer solution, and 50% PBS (–) in HBSS (–) without phenol red. After that, fluorescence signal was measured for 10 min using FDSS/μCELL (Hamamatsu Photonics). Peak numbers were evaluated using wave checker software (Hamamatsu Photonics).

### 2.9 Statistical analysis

Statistical analysis was performed using GraphPad Prism 10.4.1 software. For qRT-PCR analysis comparing two groups, a two-tailed unpaired Student’s t-test was performed. For measurement of TNF-α and IL-1β proteins comparing four groups, statistical analysis was performed using 2-way analysis of variance (ANOVA), followed by the Bonferroni’s post-hoc test. For Ca^2+^ imaging assay, statistical analysis was performed using 2-way ANOVA, followed by the Dunnett’s post hoc test.

## 3 Results

### 3.1 Microglia with branched morphology were changed into amoeboid shape by LPS stimulation in the tri-culture system

To construct a tri-culture system replicating a homeostatic state of the human brain, we firstly generated iPSC-derived astrocytes. Since it is known that astrocytes are irreversibly activated upon exposure to serum (Foo et al., 2011), we generated astrocytes from astrocyte progenitor cells (APCs) by stimulation with BMP4 and CNTF without serum. Compared to iPSCs, astrocytes showed increased levels of astrocyte marker expression (Fig. 1A). We also confirmed the expression of GFAP by immunostaining. Astrocytes exhibited a greater number of cells expressing GFAP compared to APCs (Fig. 1B). Next, a tri-culture system was established using human iPSC-derived neurons, astrocytes, and microglia. Microglia and neurons were seeded onto astrocytes, and the cells were cultured for 4-weeks. Then we confirmed a neuroinflammatory state by evaluating the morphology of microglia. Microglia are known to show branched processes in a homeostatic state and to turn into an amoeboid shape in a neuroinflammatory state (Vidal-Itriago et al., 2022). In our tri-culture system, microglia stained with IBA1, a marker of microglia, showed a morphology with branched processes (Fig. 1C). In contrast, microglia in a monoculture showed amoeboid morphology without branched processes after 4 weeks culture (Fig. 1D). These results suggested that microglia in the tri-culture system were a homeostatic state compared to the monoculture. Next, the tri-culture system was stimulated with LPS to induce a neuroinflammatory state. LPS is known to stimulate microglia specifically and to induce neuroinflammation in human cells (Tarassishin et al., 2014). The tri-culture system was treated with 100 ng/mL of LPS, and the morphology of microglia was observed at 0.5, 1, 3, 6, and 24 hr by immunofluorescent staining of IBA1. The immunostaining indicated that microglia turned into an amoeboid shape after 24 hr of LPS stimulation (Fig. 2A). In addition, the gene expression of pro-inflammatory cytokines, such as *TNF* and *IL1B*, was increased in the tri-culture system by stimulation with LPS (Fig. 2B). Conversely, the co-culture system consisting of astrocytes and neurons did not show an increase in the gene expression of pro-inflammatory cytokines upon LPS stimulation (Fig. 2B).

**Figure 1.**
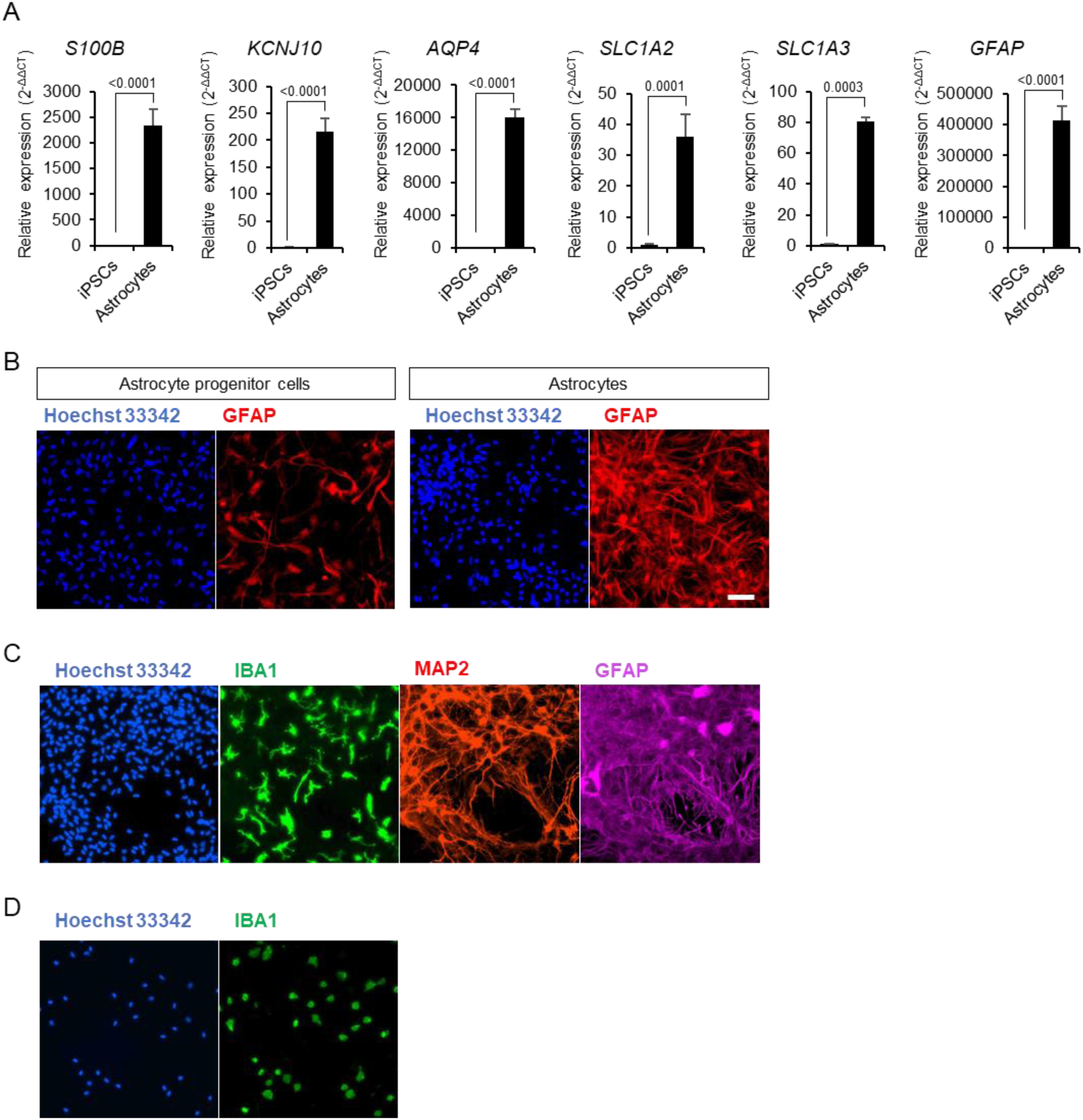
Microglia exhibited a more homeostatic morphology in the tri-culture system compared to the monoculture of microglia. (A) Quantification of the expression of astrocyte markers by qRT-PCR in iPSCs and iPSC-derived astrocytes (Data are presented as mean values and SEM. N=3.). (A-D) Fluorescence images of (B) astrocyte progenitor cells and astrocytes, (C) the tri-culture system of neurons, astrocytes and microglia, and (D) the monoculture of microglia. Scale bar: 100 μm.

**Figure 2.**
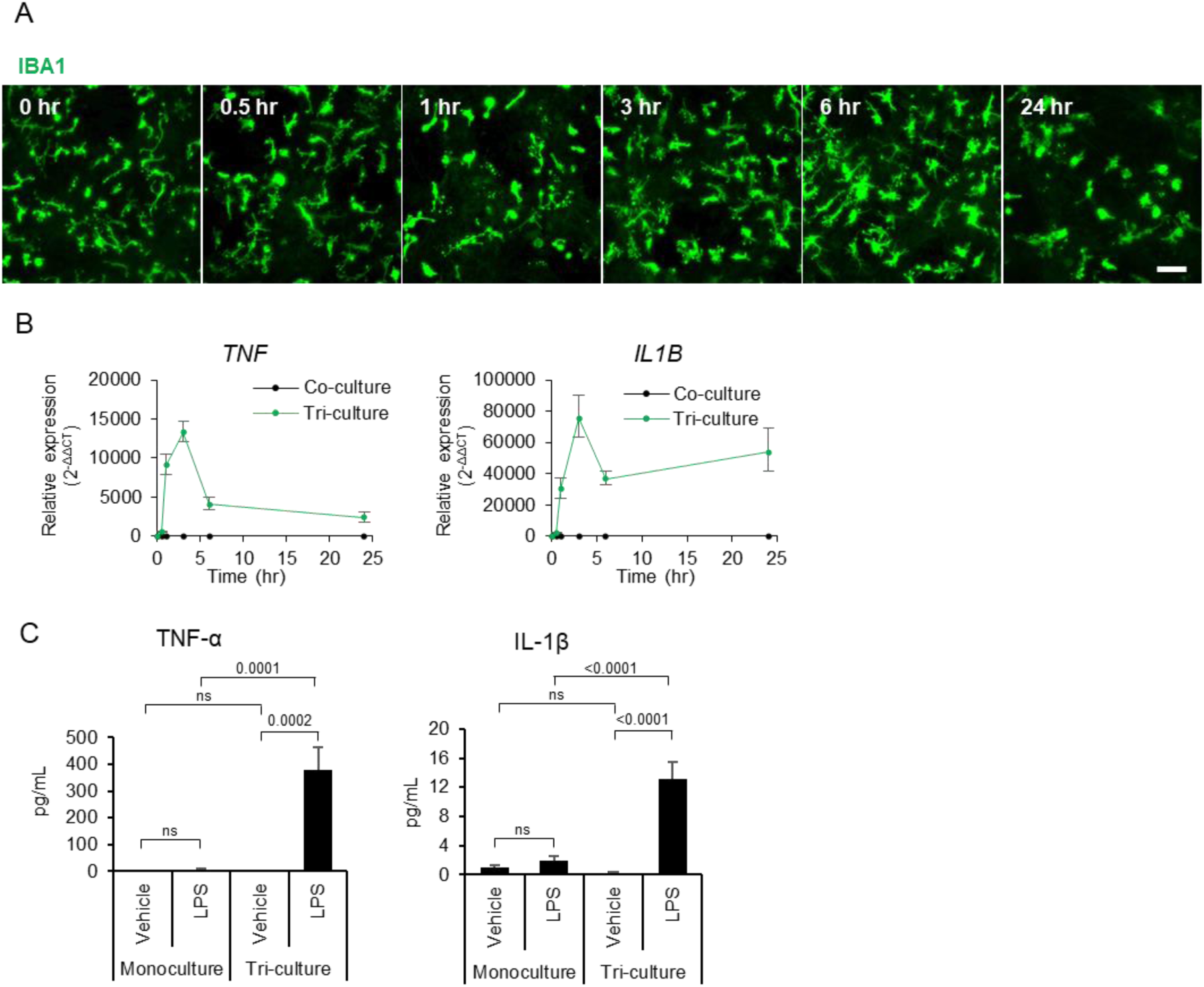
LPS stimulation in the tri-culture system induced morphological changes in microglia and the expression of pro-inflammatory cytokines. (A) Time-course fluorescent images of microglia in the tri-culture system stimulated with LPS (100 ng/mL, 24 hours). Scale bar: 100 μm. (B) Time-course analysis of gene expression for pro-inflammatory cytokines in co- and tri-culture systems by qRT-PCR at 0, 0.5, 1, 3, 6, and 24 hours after LPS stimulation (Data are presented as mean values and SEM. N=3). (C) Quantification of TNF-α and IL-1β in the culture medium following LPS stimulation (100 ng/mL, 24 hours) of the microglia monoculture and tri-culture systems (Data are presented the average of data from 3 wells and SD.)

The protein expression levels of TNF-α and IL-1β were evaluated in both the microglia monoculture and the tri-culture system (Fig. 2C). In the microglia monoculture, a slight increase in TNF-α and IL-1β levels in the medium was observed by LPS stimulation (unstimulated (vehicle): TNF-α, 1.3 pg/mL; IL-1β, 1.4 pg/mL; LPS-stimulated: TNF-α, 8.2 pg/mL; IL-1β, 2.7 pg/mL). On the other hand, in the tri-culture system, LPS stimulation induced much higher levels of TNF-α and IL-1β in the medium (unstimulated (vehicle): TNF-α, 0.7 pg/mL; IL-1β, 0.3 pg/mL; LPS-stimulated: TNF-α, 522.3 pg/mL; IL-1β, 18.1 pg/mL). These results indicate that the tri-culture system induce stronger neuroinflammatory response to LPS compared to the microglia monoculture.

### 3.2 Microglia were necessary to activate astrocytes in the tri-culture system

TLR4 is known as a receptor of LPS. Binding of LPS to TLR4 leads to the dissociation of NF-κB from IκB, allowing NF-κB translocation from the cytosol into the nucleus and the subsequent increase in the expression of pro-inflammatory cytokines, such as *TNF* (Zusso et al., 2019). Then, we evaluated the localization of NF-κB in a time-course manner following LPS stimulation to examine which cell in the tri-culture was stimulated with LPS. In the tri-culture system, nuclear translocation of NF-κB was observed in a small number of cells at 0.5 and 1 hr after LPS stimulation, whereas it was recognized in a large number of cells at 3 and 6 hr (Fig. 3A). Next, we performed immunostaining to determine in which cells nuclear translocation of NF-κB occurred. Co-staining of IBA1 and NF-κB showed that nuclear translocation of NF-κB occurred in microglia at 0.5 hours after LPS stimulation (Fig. 3B). After 3 hours of LPS stimulation, nuclear translocation of NF-κB was observed in cell populations other than microglia (Fig. 3C). Co-staining of GFAP and NF-κB showed that nuclear translocation of NF-κB occurred in astrocytes at 3 hours after LPS stimulation (Fig. 3D). On the other hand, co-localization of NeuN and NF-κB was not observed at the timepoints (Fig. 3E). These results suggest that nuclear translocation of NF-κB first occur in microglia after LPS stimulation, and subsequently in astrocytes. In contrast, the nuclear translocation of NF-κB was not confirmed at any time points in the co-culture system without microglia (Fig. 4A), suggesting that the activation of astrocytes in the tri-culture system is not directly induced by LPS stimulation but by activated microglia and that microglia are necessary to activate astrocytes. Next, we evaluated the population of microglia required to activate astrocytes. The localization of NF-κB was assessed after LPS stimulation in tri-cultures with microglial ratio of 0.1%. Despite the low proportion of microglia in the tri-culture system, LPS stimulation induced the nuclear translocation of NF-κB in microglia followed by astrocytes (Fig. 4B). Thus, the small amount of microglia, such as 0.1% of cell population, was enough to activate astrocytes.

**Figure 3.**
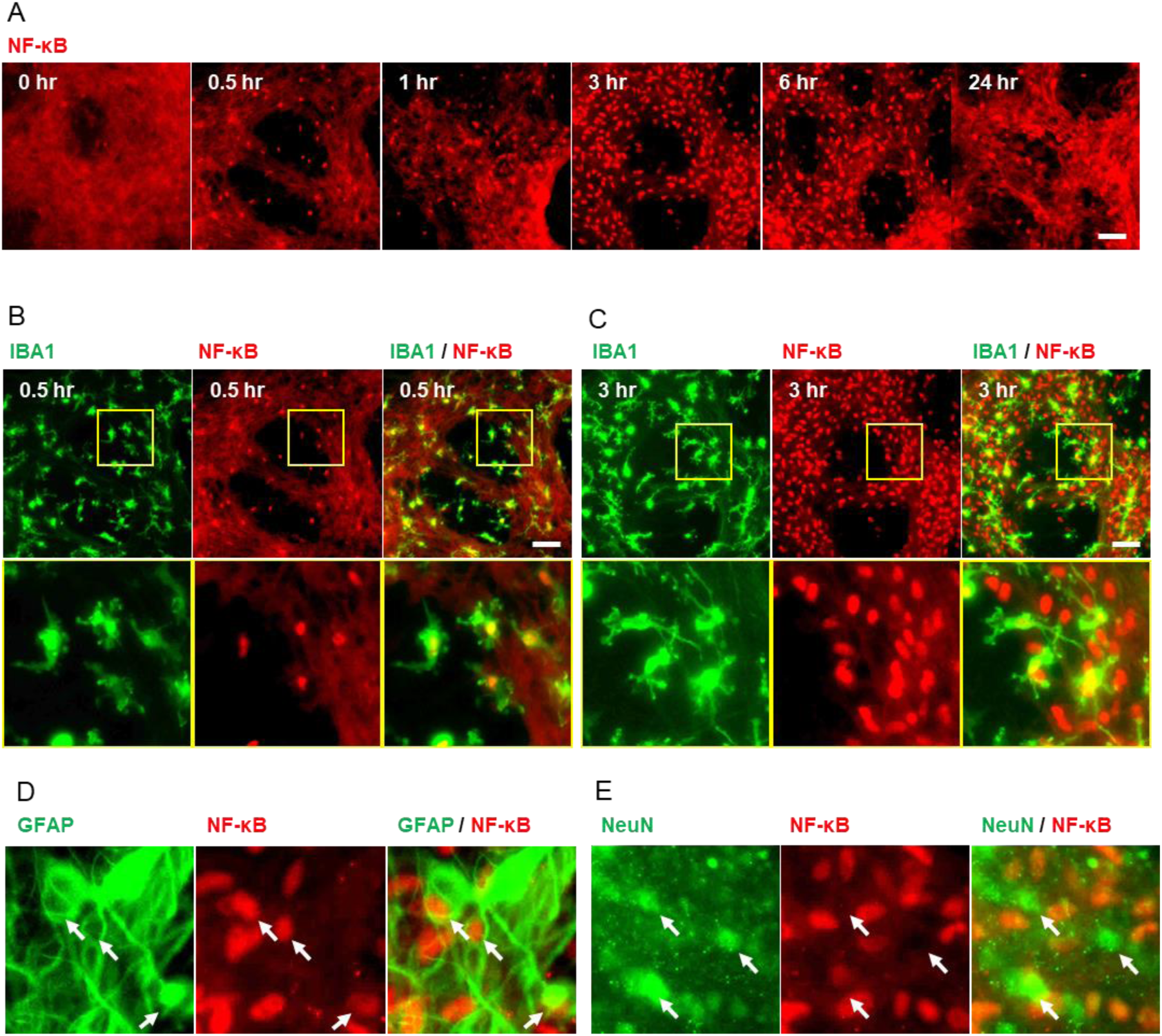
LPS stimulation in the tri-culture system first activated microglia, followed by the activation of astrocytes. (A) Time-course fluorescent images showing localization changes of NF-κB in the tri-culture system stimulated with LPS (100 ng/mL, 24 hours). (B, C) Fluorescent images of NF-κB and microglial marker IBA1 in the tri-culture system at 0.5 and 3 hours after LPS stimulation. (D) Fluorescent images of NF-κB and astrocytic marker GFAP in the tri-culture system at 3 hours after LPS stimulation. (E) Fluorescent images of NF-κB and neuronal marker NeuN in the tri-culture system at 3 hours after LPS stimulation. Scale bar: 100 μm.

**Figure 4.**
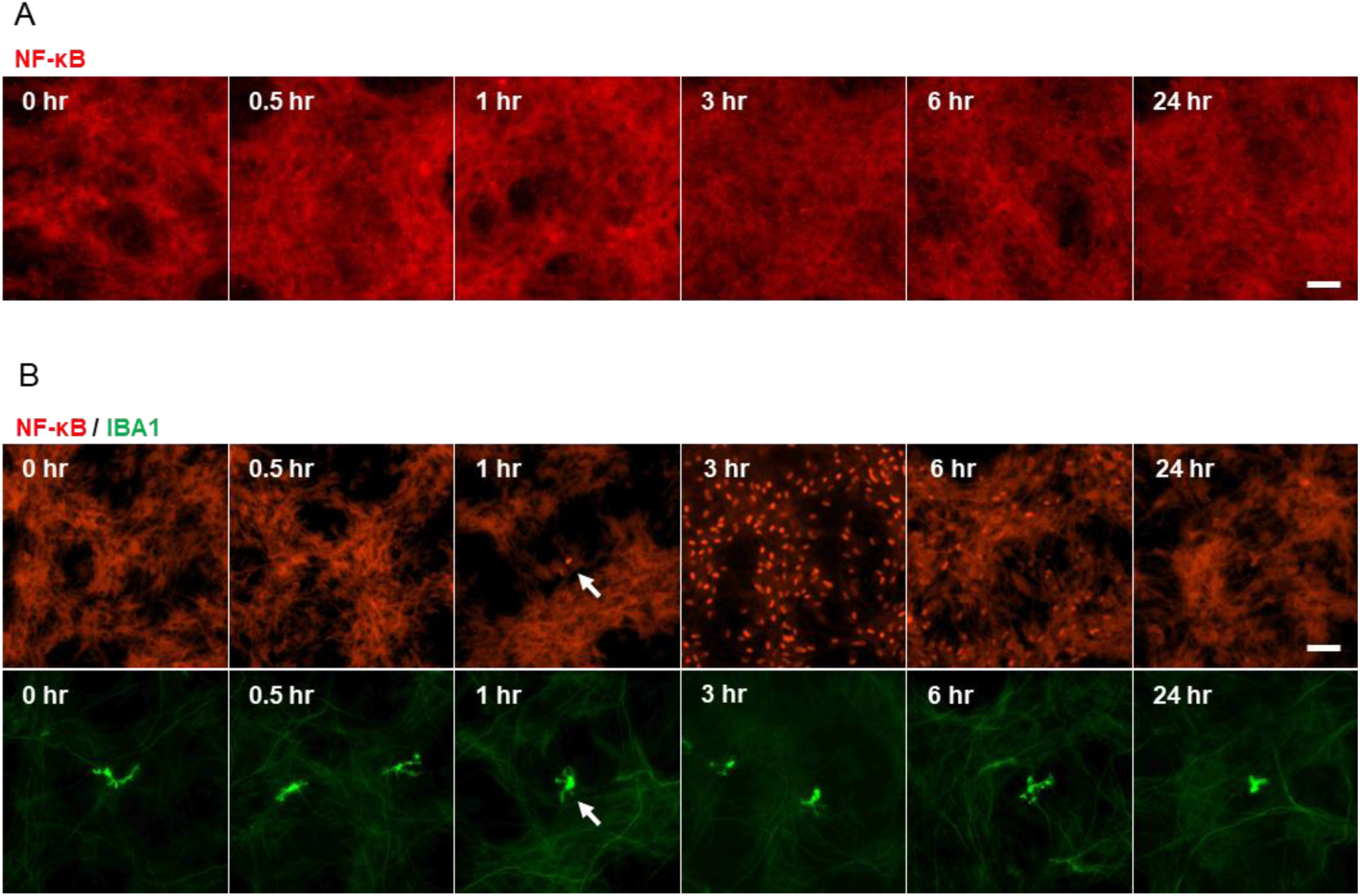
Microglia were necessary to activate astrocytes in the tri-culture system. (A) Time-course fluorescent images of NF-κB in the neuron-astrocyte co-culture system stimulated with LPS (100 ng/mL, 24 hours). (B) Time-course fluorescent images showing the localization changes of NF-κB in the tri-culture system containing 0.1% of microglia after LPS stimulation (100 ng/mL, 24 hours). Scale bar: 100 μm.

### 3.3 LPS-stimulated microglia transduced signal to astrocytes through TNF-α

We further investigated the molecular mechanism of nuclear translocation of NF-κB in astrocytes induced by LPS-stimulated microglia. It is known that activated microglia increase the *TNF* expression and that NF-κB nuclear translocation occurs in downstream of TNF-α binding to TNF receptor (Bouwmeester et al., 2004). Then, we examined whether TNF-α signal was involved in the NF-κB translocation in astrocytes following LPS stimulation in the tri-culture system. R-7050 was used to inhibit the binding of TNF-α to TNF receptors. R-7050 was applied simultaneously with LPS stimulation, and the NF-κB localization was assessed in the tri-culture system. The nuclear translocation of NF-κB was confirmed in a small number of cells at 1, 3, and 6 hr after LPS stimulation (Fig. 5A). Immunofluorescent staining was conducted to identify the cell types showing the nuclear translocation of NF-κB, and it was observed that all cells displaying accumulation of NF-κB in the nuclear were IBA1 positive (Fig. 5B). These findings indicated that the TNF receptor inhibitor suppressed the astrocytes activation in the LPS-stimulated tri-culture system. LPS stimulation in the tri-culture system led to the nuclear translocation of NF-κB and an upregulation of *TNF* expression in microglia. Subsequently astrocytes were activated by the TNF-α molecules released from microglia, followed by the nuclear translocation of NF-κB.

**Figure 5.**
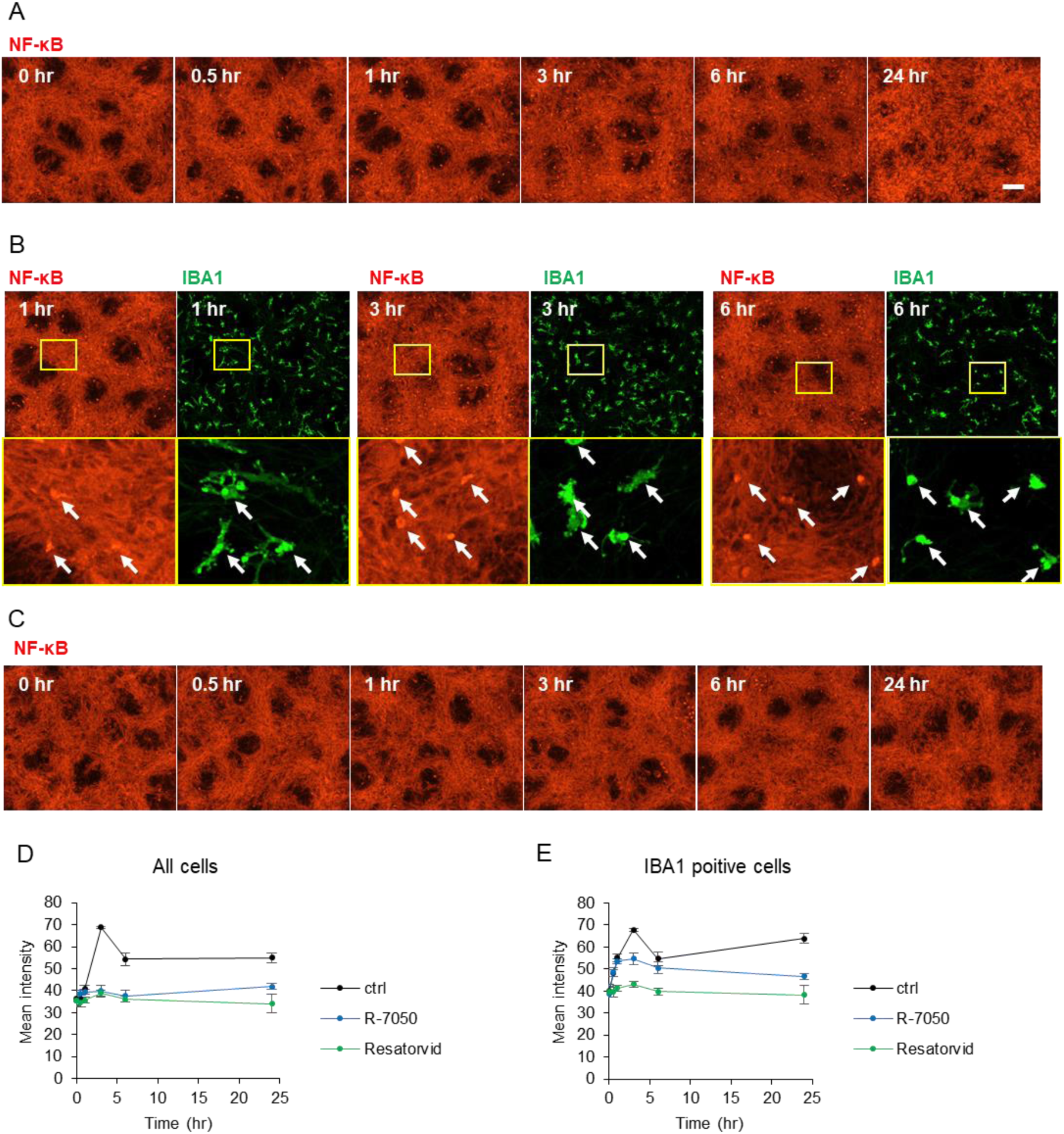
TNF receptor inhibitor suppressed astrocyte activation by activated microglia after LPS stimulation. (A) Time-course fluorescent images showing the localization of NF-κB in the tri-culture system stimulated with LPS (100 ng/mL) and R-7050 (10 μM), a TNF receptor inhibitor, for 24 hours. (B) Fluorescent images of NF-κB and microglial marker IBA1 in the tri-culture system at 1, 3 and 6 hours after LPS stimulation. (C) Time-course fluorescent images of NF-κB in the tri-culture system stimulated with LPS (100 ng/mL) and resatorvid (1 μM), a TLR4 inhibitor, for 24 hours. Scale bar: 100 μm. (D, E) Quantitative data of fluorescence intensity for NF-κB signaling in the nuclear regions of (D) all cells or (E) IBA1 positive cells (Data are presented the average of data from four fields and SD).

In addition, we evaluated the effects of TLR4 inhibitor on LPS-induced microglial activation. Treatment of the TLR4 inhibitor Resatorvid with LPS inhibited NF-κB nuclear translocation at all time points (Fig. 5C). This indicates that microglia are stimulated with LPS through TLR4. Furthermore, quantification of NF-κB fluorescence intensity in the nuclear regions of all cells revealed a slight increase in intensity at 0.5 and 1 hour in the vehicle group, with a marked increase observed at 3 and 6 hours (Fig. 5D). These results align with the finding that the nuclear translocation of NF-κB occurs only in microglia at 0.5 and 1 hour and that the nuclear translocation is also observed in astrocytes at 3 and 6 hours. The treatments with R-7050 and Resatorvid clearly inhibited nuclear translocation at 3 and 6 hours. Next, we quantified NF-κB fluorescence intensity limited to the nuclear regions of IBA1-positive cells. It demonstrated that R-7050 did not inhibit the nuclear translocation at the 0.5 and 1 hour, whereas significant inhibition was observed at the 3 and 6-hour points (Fig. 5E). These data indicate that R-7050 does not inhibit LPS-induced microglial activation, but rather suppresses the activation of astrocytes by TNF-α released from the activated microglia. On the other hand, Resatorvid inhibited the nuclear translocation of NF-κB at all time points, indicating the inhibition of LPS-induced microglial activation by Resatorvid (Fig. 5E).

### 3.4 The activated astrocytes enhanced neuronal excitation

It has been reported that neuronal excitability is enhanced in a neuroinflammatory state (Rodgers et al., 2009). To confirm the neuronal activity in our tri-culture system under a condition of neuroinflammation, we conducted a calcium imaging assay. Calcium oscillation, which is synchronized repetitive calcium influxes, was quantified to evaluate neuronal activities. The number of peaks per minute (beats per minute: BPM) did not change in the neuron-astrocyte co-culture system treated with LPS for 24 hours, whereas an increase in BPM was observed in the tri-culture system with LPS stimulation (Fig. 6A). These results suggest that neuronal excitation is enhanced in the tri-culture system under a condition of neuroinflammation. In our try-culture system, microglia activated astrocytes through TNF-α after LPS stimulation. We examined the involvement of TNF-α-stimulated astrocytes in neuronal excitation. In both neuron-astrocyte co-culture and tri-culture systems, TNF-α stimulation resulted in enhanced neuronal excitation (Fig. 6A). It is known that the binding of TNF-α to TNF receptor 1 leads to the dissociation of NF-κB from IκB, allowing NF-κB translocation into the nucleus. To determine which cells, either neurons or astrocytes, are stimulated by TNF-α, we examined the localization of NF-κB. In the neuron-astrocyte co-culture system, GFAP positive cells showed the nuclear translocation of NF-κB after 3 hours of TNF-α stimulation (Fig. 6B). In contrast, NeuN positive cells did not exhibit nuclear translocation of NF-κB after TNF-α stimulation (Fig. 6C). Additionally, we confirmed the localization of NF-κB in each monoculture system. In monoculture of astrocytes, the translocation of NF-κB was confirmed after 3 hours of TNF-α stimulation (Fig. 6D). On the other hand, accumulation of NF-κB was not observed in monoculture of neurons (Fig. 6E). These results suggest that TNF-α stimulates astrocytes rather than neurons, and that neuronal excitation is enhanced through the activation of astrocytes. In our study, the cellular interaction in a tri-culture system under the condition of a neuroinflammatory state was occurred as follows: 1) microglia were activated and released TNF-α by LPS stimulation, 2) activated microglia subsequently activated astrocytes by releasing TNF-α, 3) the activated astrocytes enhanced neuronal excitation (Fig. 7).

**Figure 6.**
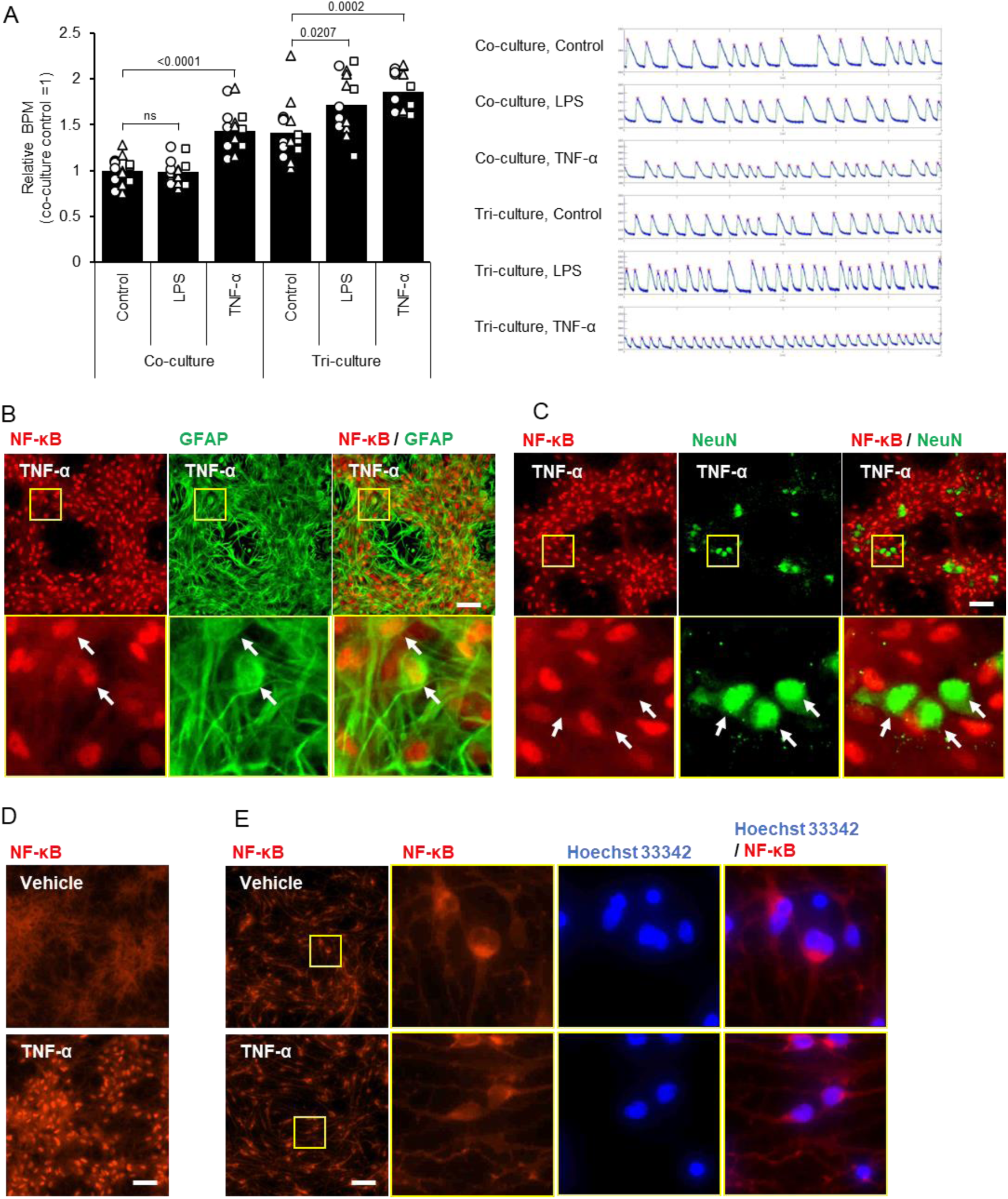
**Neuronal activity was increased by activated astrocytes.**(A) Neuronal activity measured by calcium imaging in the co- and tri-culture systems stimulated for 24 hours with LPS (100 ng/mL) or TNF-α (7.5 ng/mL). (Data are presented the average of 3 independent experiments with SEM. The data from each experiment are represented by indibidual markers (circles, triangles, and squares) and include measurements from 3-6 wells. All values relative to control in co-culture). (B, C) Fluorescent images of NF-κB and astrocyte marker GFAP or neuronal marker NeuN in the co-culture system at 3 hours after TNF-α (7.5 ng/mL) stimulation. (D, E) Fluorescent images of NF-κB in the monoculture of (D) astrocytes or (E) neurons at 3 hours after TNF-α (7.5 ng/mL) stimulation. Scale bar: 100 μm.

**Figure 7.**
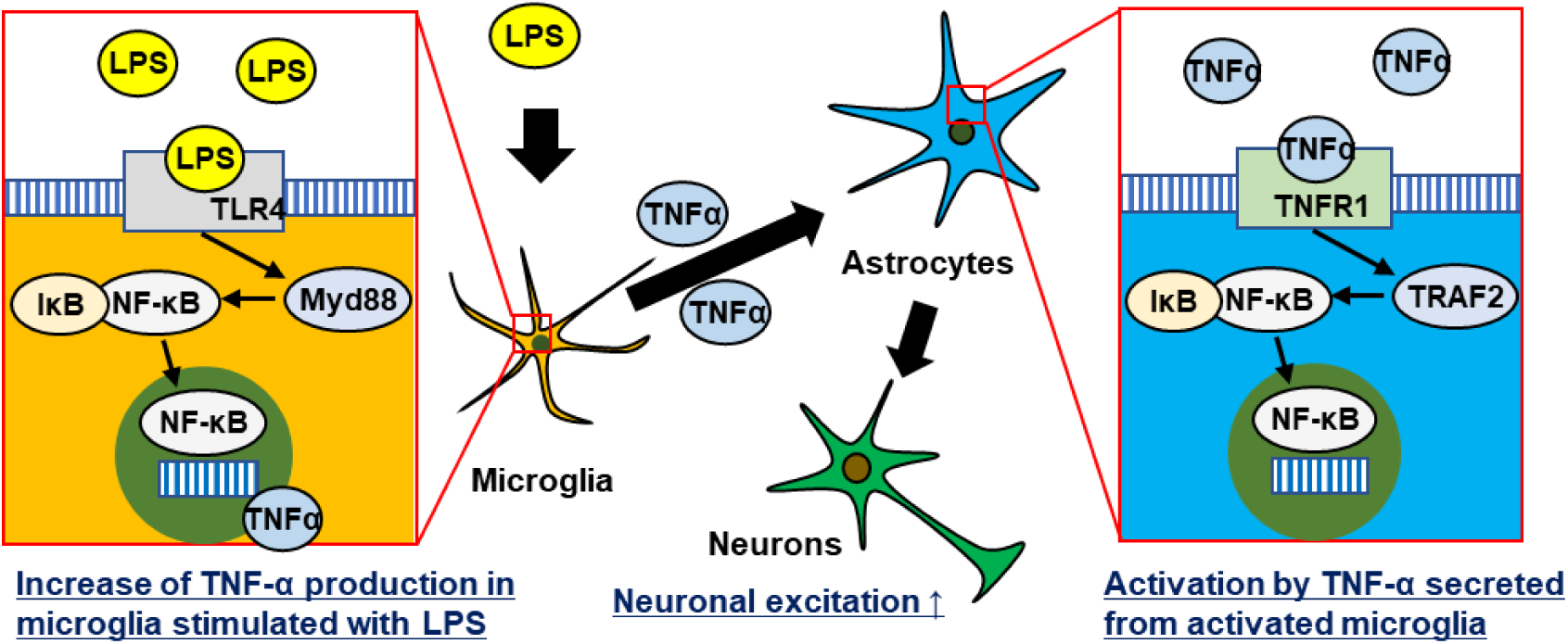
**Schematic image of cellular interaction of the tri-culture system stimulated with LPS.**The tri-culture system revealed that microglia were first activated by LPS, accompanied by the nuclear translocation of NF-κB and the release of TNF-α. Subsequently, TNF-α activated astrocytes, leading to increased neuronal excitation.

## 4 Discussion

In this study, the tri-culture system successfully reproduced a transition from a homeostatic state to a neuroinflammatory state, uncovering the cellular interaction among microglia, astrocytes, and neurons during neuroinflammation. The presence of microglia is essential to induce neuroinflammation in response to LPS, while the presence of astrocytes and neurons is necessary to amplify the neuroinflammatory response. Through visualization of NF-κB nuclear translocation, we observed that microglia were activated first, followed by astrocyte activation.

Microglia in the tri-culture system showed morphology with branched processes, while microglia in monoculture showed an amoeboid shape, suggesting that the tri-culture system more closely resembled a homeostatic state. Furthermore, the tri-culture system showed lower levels of TNF-α and IL-1β expression compared to the monoculture, supporting the idea that microglia in the tri-culture system represent a more homeostatic state. Our results are also in line with a previous study that reporting microglia exhibiting a state of homeostasis when co-cultured with neurons and astrocytes (Baxter et al., 2021). The report highlights the crucial role of TGF-β2 signaling in maintaining microglial homeostasis. In addition, BDNF and GDNF are also known to inhibit microglial activation (Wu et al., 2020; Rocha et al., 2012). It is postulated that the release of various factors from neurons and astrocytes influences the state of microglia in the tri-culture system.

The tri-culture system exhibited a more pronounced increase in cytokine levels compared to the monoculture system upon LPS stimulation. Maintaining a homeostatic state may be important for responding to stimuli. In addition, the increased cytokine levels in the tri-culture system implies that the presence of astrocytes and neurons is essential for amplifying the inflammatory response. Our study demonstrated astrocytes activation by microglia stimulated with LPS. Furthermore, it is known that activated astrocytes can activate microglia (Tanuma et al., 2006). In the tri-culture system, astrocytes and microglia may have contributed to a more robust inflammatory response through bi-directional cellular interactions.

In our study, the nuclear translocation of NF-κB in astrocytes and neurons was not observed in the neuron-astrocyte co-culture system after LPS stimulation. However, the nuclear translocation of NF-κB in astrocytes was observed in the tri-culture system stimulated by LPS. These results suggest that the presence of microglia is essential to respond to LPS stimulation. It has been reported that human astrocytes do not respond to LPS stimulation (Tarassishin et al., 2014), while there is a report that human primary astrocytes can respond to LPS (Hao et al., 2001). Rodent primary astrocytes have also been reported to respond to LPS (Lieberman et al., 1989; Galindo et al., 2011). Our study demonstrated that even a small population of microglia, such as 0.1% of the cell population, was sufficient to induce astrocyte activation. The activation of primary astrocytes by LPS reported in the literature could be attributed to the contamination of a small amount of microglia. However, we cannot exclude the possibility that the differences of astrocyte responses are due to heterogeneity among astrocytes based on the brain region or species differences between human and rodents. These variations have been previously reported and cannot be excluded as potential factors influencing astrocyte responses (Batiuk et al., 2020; Zhang et al., 2016).

It has been reported that administration of LPS to rodents induces seizures (Rodgers et al., 2009). In our study, enhancement of neuronal activity was confirmed by LPS stimulation in the tri-culture system. Furthermore, we indicated that neuronal activity was enhanced through the activation of astrocytes, which resulted from TNF-α released by activated microglia. Astrocytes are known to regulate neuronal activity through glutamate (Blanco-Suárez et al., 2017). It is also known that the activated astrocytes lead to reduced expression of glutamate transporters (Leventoux et al., 2020; Tilleux et al., 2008). Thus, it is possible that the alteration of neuronal activity in the tri-culture system after LPS stimulation may be associated with changes in the glutamate concentration in the medium or in the local environment.

In our study, we conducted calcium imaging assay to evaluate neuronal activity. Calcium oscillations, characterized by synchronized repetitive calcium influxes, are observed following the formation of a neuronal network (Kuijlaars et al., 2016). In a tri-culture system using rodent cells, a previous study has reported the evaluation of synchronized neural excitability (Phadke et al., 2022). However, there have been few reports evaluating the neuronal network activity in a tri-culture system using human iPSC-derived cells. In our tri-culture system, calcium oscillation was confirmed, suggesting that the neuronal cells are mature and formation of a neuronal network and that they are overcoming the immaturity issues of iPSC-derived neuronal cells (Ohara et al., 2015; Odawara et al., 2016). The tri-culture system is valuable for research on mental disorders and pain, where the involvement of glial cells in neural activity is noted. For example, trisomy 21, which is known to cause Down syndrome, has been reported to activate microglia with morphological changes, leading to alterations in neural activity (Pinto et al., 2020; Zheng et al., 2021). In neuropathic pain, it has been reported that the activation of microglia is involved in the overactivity of neurons (Kohno et al., 2022). Our tri-culture model provides a valuable tool for investigating the pathogenesis of these diseases and uncovering the role of glial cells in these diseases.

In conclusion, we achieved to generate microglia in a more homeostatic state compared to the monoculture condition by utilizing the tri-culture system. We revealed the cellular interaction in which astrocytes are activated by TNF-α released from microglia under a neuroinflammation state, leading to enhanced neuronal activity. The utilization of our tri-culture system provides a valuable tool for evaluating cellular interactions among neurons, astrocytes, and microglia, thereby facilitating a better understanding of neuroinflammation and neurodegenerative diseases.

## Abbreviations

iPSCs: induced pluripotent stem cells
LPS: lipopolysaccharide
NF-κB: nuclear factor kappa
B TNF-α: tumor necrosis factor-alpha
ANOVA: analysis of variance
APCs: astrocyte progenitor cells
BPM: beats per minute

## Author contributions

H.Kobayashi, H.Kato, and S.Endoh-Yamagami contributed conceptualization. H.Kobayashi and M.Taniguchi performed the experiments in this study and analyzed data. H.Kobayashi, and S.Endoh-Yamagami wrote the original manuscript. S.Endoh-Yamagami supervised the project. All authors reviewed the manuscript and approved the final manuscript.

## Acknowledgments

The authors would like to thank Kazuhiko Nakata and Ayako Sakai for their assistance in the experiments. The authors also would like to thank Akira Nabetani and colleagues of Fujifilm corporation for helpful discussions and support.

